# Sofosbuvir and its tri-phosphate metabolite inhibit the RNA-dependent RNA polymerase activity of non-Structural protein 5 from the Kyasanur forest disease virus

**DOI:** 10.1101/2022.06.29.498065

**Authors:** Mansi Malik, Parvathy Vijayan, Deepak K Jagannath, Rakesh K Mishra, Anirudha Lakshminarasimhan

## Abstract

Kyasanur forest disease is a neglected zoonotic disease caused by a single-stranded RNA-based flavivirus, the incidence of which was first recorded in 1957 in the Southern part of India. Kyasanur forest disease virus is transmitted to monkeys and humans through the infected tick bite of *Haemophysalis spinigera*. Kyasanur forest disease is a febrile illness, which in severe cases, results in neurological complications leading to mortality. The current treatment regimens are symptomatic and supportive, and no targeted therapies are available for this disease. In this study, we evaluated the ability of FDA-approved drugs sofosbuvir (and its active metabolite) and Dasabuvir to inhibit the RNA-dependent RNA polymerase activity of NS5 protein from the Kyasanur forest disease virus. NS5 protein containing the N-terminal methyl transferase domain and C-terminal RNA-dependent RNA polymerase domain was expressed in *Escherichia coli*, and RNA-dependent RNA polymerase activity was demonstrated with the purified protein. The RNA-dependent RNA polymerase assay conditions were optimized, followed by the determination of apparent K_m,ATP_ to validate the enzyme preparation. Half maximal-inhibitory concentrations against RNA-dependent RNA polymerase activity were determined for Sofosbuvir and its active metabolite. Dasabuvir did not show detectable inhibition with the tested conditions. This is the first demonstration of the inhibition of RNA-dependent RNA polymerase activity of NS5 protein from the Kyasanur forest disease virus with small molecule inhibitors. These initial findings can potentially facilitate the discovery and development of targeted therapies for treating Kyasanur forest disease.

## Introduction

Kyasanur forest disease (KFD) is an infectious viral hemorrhagic disease primarily infecting human and non-human primates [1]. Mortality in monkeys, especially among the black-faced langurs (*Presbytis entellus*) *a*nd red-faced bonnet monkeys (*Macaca radiata*), has been reported during multiple outbreaks of this disease [2]. In addition, birds, squirrels, gerbils, mice, rats, and bats also function as hosts for the virus [3]. The hard-bodied tick, *Haemaphysalis spinigera*, is the primary vector for the Kyasanur forest disease virus (KFDV) [4]. KFDV was also isolated from various genera of ticks, such as *Ixodes, Argas, Ornithodoros, Hyalomma, Dermacentor, and Rhipicephalus* [3]. Clinical symptoms of KFD include high-grade fever, chills, headache, Conjunctivital inflammation, vomiting, abdominal pain, diarrhea, maculopapular eruptions, muscle tenderness, nausea, and persistent cough [5]. In extreme cases, neurological and hemorrhagic manifestations are observed, with an estimated fatality of ~ 3-5% [6]. A microchip-based point-of-care test (Truenat KFD), developed by ICMR-NIV and Molbio Diagnostics, Goa, is currently the gold standard for diagnosis in India [7]. Formalin inactivated tissue-culture vaccine is available for KFD, administered in two doses to individuals aged 7–65 years at one-month intervals [8]. However, the immunity conferred by this vaccine is short-lived because of which booster doses are recommended after 6–9 months of primary vaccination for five consecutive years [8]. Current treatment regimens for KFD are limited to providing symptomatic relief and management [6]. There are no direct-acting antiviral drugs available against KFD. Failure of optimal coverage provided by the vaccine, with the absence of antiviral interventions against KFDV, makes it a public health challenge, necessitating attention toward developing second-generation vaccines and targeted therapies.

KFDV is a spherical (~40-65 nm in size) virus containing an icosahedral nucleocapsid enclosing a positive-sense RNA of 10,774 bases [9]. Kyasanur forest disease virus genome is closely related to Alkhurma Virus from Saudi Arabia, Omsk hemorrhagic fever virus from Siberia, and Nanjianyin Virus from China [10–12]. The RNA strand codes for a polyprotein comprising three structural (C, E & M) and seven non-structural proteins (NS1, NS2a, 2b, NS3, NS4a, 4b, and NS5 [13]. NS5 belonging to the Flaviviridae family are bi-functional proteins comprising of an N-terminal methyl transferase domain (MTase) and a C-terminal RNA-dependent RNA polymerase domain (RdRp) separated by a linker [14, 15](Fig. 1A). MTase domain methylates N-7 of guanine and 2’-OH ribose of viral RNA cap, in the presence of S-adenosyl methionine as a co-factor. N-7 methylation is critical for enhancing the efficiency of viral translation, and 2’-O methylation is essential for the subversion of the innate immune response during viral infection [15]. RdRp domain of flaviviruses catalyzes the de novo template-dependent formation of phosphodiester bonds in the presence of nucleoside triphosphate (NTP) and divalent metal ions. The canonical structure of the Flaviviridae RdRp domain consists of an encircled right hand containing three subdomains - thumb, palm, and fingers folded, surrounding the active site. RdRps consist of channels, the entry of which is lined with positively charged residues, facilitating NTPs and template RNA entry into the active site [14]. The absence of proofreading exonuclease activity in RdRps may lead to the incorporation of mutations and, consequently, the selection of variants of viruses [16]. Hence, RdRp of NS5 plays a crucial role in viral genome replication and can serve as a potential antiviral drug target [17]. Nucleoside and non-nucleoside inhibitors against NS5 RdRp from different viruses belonging to the Flaviviridae family have been identified [18–25].

**Fig. 1.**
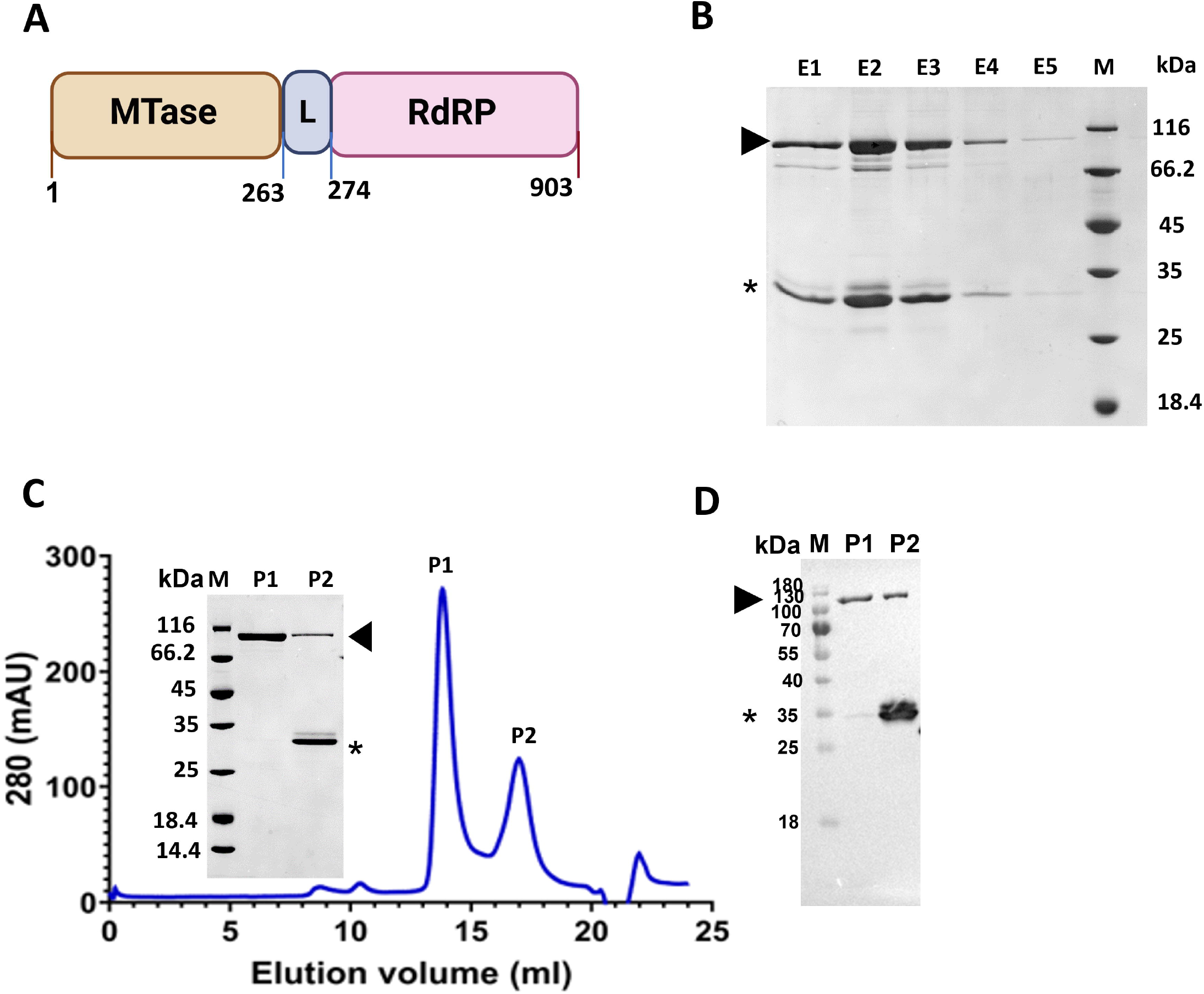
NS5 protein of Kyasanur forest disease virus. A. Schematic representation of NS5 protein domains - N-terminal methyl transferase (MTase – 1 to 263) and C-terminal RNA dependent RNA polymerase (RdRp – 274 to 903). Linker (L – 264 to 273 amino acids) residues connecting the two domains are not conserved among flaviviruses (35). B. Coomassie-stained SDS-PAGE gel (12%) containing Ni-NTA affinity column elutes. Lanes containing a prominent band of the expected molecular weight of 104 kDa, corresponding to 5 to 897 amino acids of NS5, are seen in the elutes (indicated by an arrow). Also seen is a lower molecular weight band between the molecular weight markers - 35 kDa and 25 kDa (*). C. Superdex increase 200 10/300 GL fractionation of affinity purified fraction of NS5 protein. Peaks P1 and P2 correspond to NS5 and the lower molecular weight protein (*). Coomassie-stained SDS-PAGE gel is on the left panel. D. anti-Histidine tagged antibody probed western blot of P1 and P2 fractions. The prominent bands in P1 (NS5 protein indicated with arrowhead) and P2 (lower molecular weight contaminant band marked with asterisks) were recognized by an anti-His antibody.

Sofosbuvir is an FDA-approved drug currently in use for the treatment of Hepatitis C virus (HCV) infection [26, 27]. In this study, we recombinantly expressed KFDV NS5 protein and characterized the purified protein to identify a potential inhibitor of RdRp activity. Sofosbuvir (Pro-drug) and its metabolite were demonstrated to inhibit the RdRp activity of the KFDV NS5 protein.

## Materials and Methods

### Expression and purification of NS5 from KFDV

Codon-optimized gene corresponding to 5 to 897 amino acids of KFDV NS5 with an N-terminal Hexa-histidine tag was synthesized (Genscript USA) and sub-cloned into pET-28a (+) vector between NdeI and HindIII restriction sites. Transformants of Rosetta2(DE3) pLysS containing pET28a – KFDV NS5 were grown to an OD of 2.0 and induced with 0.5 mM IPTG for 16 hours at 18°C. Induced cells were centrifuged at 5000 g for 10 minutes at 4°C, and the pellet was resuspended in a lysis Buffer containing 20 mM Na_2_HPO_4_, 0.5 M NaCl, 50 mM L-Arginine, 10 mM MgSO_4_, 5 mM β-mercaptoethanol, 5 mM Imidazole, 1 mM PMSF, 10% Glycerol, pH 7 [28]. The cell suspension was sonicated, and the lysate was centrifuged at 20000 g for 60 minutes at 4°C. The clarified lysate was filtered with Durapore PVDF 0.22 μm membrane and loaded onto a Ni-NTA affinity column pre-equilibrated with 20 mM Na_2_HPO_4_, 0.5 M NaCl, 10 % Glycerol pH 7.0. The column was washed with equilibration buffer containing 50 mM Imidazole and bound protein eluted with 20 mM Na_2_HPO_4_, 0.5 M NaCl, 400 mM Imidazole, 10% Glycerol pH 7.0. Eluates containing a prominent band corresponding to the expected molecular weight of ~104 kDa were injected into Superdex 200 Increase 10/300 GL column pre-equilibrated with 50 mM Tris-HCl pH 8.0, 150 mM NaCl, 5 mM MgCl_2_, 1 mM DTT, 5% Glycerol, 0.05% Tween 20. The purified protein was quantified using Bradford assay and densitometry analysis. Purified fractions were transferred onto the PVDF membrane and probed with mouse anti-His primary antibody (1:3000) and anti-mouse IgG HRP—conjugate (1:6000). Gel peptic digest and LC-MS/MS analysis were performed for the gel filtration fractions to ascertain the identity of the NS5 protein.

### RNA-dependent RNA polymerase activity of purified NS5

RNA-dependent RNA polymerase (RdRp) activity of KFDV NS5 protein was determined by employing a fluorometric endpoint assay based on the intercalating property of a fluorescent dye, SYTO 9, which binds to double-stranded (ds) RNA and not to single-stranded (ss) RNA [29]. 250 nM of purified NS5 protein was added to a standard reaction mix comprising 50 mM HEPES pH 7.0, 2.5 mM MnCl_2_, 500 μM ATP, 20 μg/mL poly(U), and 0.1mg/mL BSA, and the reaction mix was incubated at 30°C for 60 minutes in 96 well flat-bottom black plates. The reaction was quenched by adding 25 mM EDTA, following which 0.25 μM SYTO 9 was added and further incubated at room temperature for 7 minutes. Fluorescence was recorded in a plate reader (Infinite 200 Pro M Plex, TECAN, Switzerland) using excitation and emission filters at 485 nm and 520 nm, respectively. The background fluorescence levels were subtracted from the samples using a protein blank control reaction. The effect of pH, temperature, NaCl, and glycerol concentrations on RdRp activity was determined to identify the optimal conditions using the purified KFDV NS5 protein [30]. Buffers with pH ranging from 5.0 to 9.0 (Citrate pH 5.0, Tricine pH 6.0, HEPES pH 7.0, Tris-HCl PH 7.5, Tris-HCl pH 8.0, and Bicine pH 9.0) were assayed. NS5 protein was incubated at different temperatures of 20°C, 25°C, 30°C, 37°C, and 42°C, different NaCl concentrations (5 mM, 25 mM, 50 mM, 75 mM, 100 mM, 300 mM, and 500 mM) and varying percentages of glycerol (5% v/v, 10% v/v, 20% v/v, 30% v/v, 40% v/v, 50% v/v). The mean values were computed from the experiments with standard deviation.

NS5 protein was titrated against a fixed concentration of 500 μM ATP to arrive at the protein concentration of 250 nM to determine apparent K_m,ATP_. At concentrations ranging from 50 μM to 500 μM, ATP was titrated with 250 nM of purified NS5 protein and 20 μg/mL of poly(U). Mean values were computed with standard deviations (n = 3), and K_m_ was calculated using GraphPad Prism, 9.3.0.

### Determination of inhibitory concentrations for sofosbuvir, its metabolite, and dasabuvir

Sofosbuvir, Sofosbuvir triphosphate, and Dasabuvir (MedChemExpress, Monmouth Junction, NJ) were dissolved in DMSO and diluted freshly for the experiment. The inhibitory effect of Sofosbuvir, sofosbuvir triphosphate, and Dasabuvir was determined at different concentrations ranging from 5 μM to 20 μM. The assay was performed in presence of 250 nM of NS5 protein, 20 μg/mL poly(U) and 500 μM ATP in the reaction buffer containing 50 mM HEPES pH 7.0, 5 mM NaCl, 10% glycerol, 0.1 mg/ml BSA. IC_50_ values were determined using a non-linear regression model with GraphPad Prism, 9.3.0. The mean values were computed with standard deviation (n = 3).

## Results

### KFDV NS5 was expressed in bacteria and purified with affinity column and size exclusion chromatography

Multiple sequence alignment of KFDV polyprotein (Uniprot id: D7RF80) with NS5 proteins from tick-borne encephalitis virus (TBEV), Yellow fever virus (YEFV), Zika virus (ZKV), Japanese encephalitis virus (JAEV), West Nile virus (WNV) and Dengue virus serotype 2 (DENV2), led to the identification of KFDV NS5 protein, i.e., between amino acids, 2514 to 3416 in the polyprotein sequence. NS5 protein is widely conserved across the Flaviviridae family (50 – 70% identity/similarity). Residues 5 to 897 of NS5 protein containing the MTase and RdRp domain (Fig. 1A) were expressed with an N-terminal Hexa histidine tag in bacteria and purified with Ni-NTA affinity chromatography (Fig. 1B). A lower molecular weight contaminant band (Fig. 1) was observed along with the band corresponding to the expected molecular weight of NS5 (~104 kDa), on the Coomassie blue stained SDS-PAGE gel. Anti-His antibody also recognized the lower molecular weight contaminant band with western blotting, suggesting it to be an N-terminal degradation product. It was confirmed with amino acid sequencing of tryptic digests with LC-MS/MS. The sequence corresponded to 5 to 282 amino acids of the NS5 protein (Data not shown). This histidine-tagged N-terminal fragment co-eluting with NS5 protein in affinity chromatography was removed with superdex 200 increase 10/300 GL size exclusion column chromatography (Peak P2 from Fig. 1C) to the extent that it was not observed in a Coomassie-stained SDS-PAGE gel. The identity of the NS5 protein - Peak, P1 corresponding to the expected molecular weight of 104 kDa, was confirmed with LC-MS/MS analysis and western blot analysis using an anti-His antibody (Fig. 1D). The yield of NS5 protein after size exclusion chromatography (P1 fraction) was 1 mg/liter of bacterial culture.

### Apparent K_m,ATP_ for NS5 RDRP from KFDV is comparable to that determined for other viruses

KFDV NS5 construct containing both the MTase and RdRp domains was used in this study to rule out any effect of the MTase domain on RdRp activity. The interplay of both the RdRp and MT domains was earlier demonstrated for Dengue and Zika NS5 [31–33]. Compared to 5% glycerol, KFDV NS5 showed an increase in RdRp activity at 10% glycerol, which significantly reduced at 20% (Fig 2). Inhibition of RdRp activity of KFDV NS5 was observed at lower concentration ranges of NaCl consistent with an earlier report from Usutu virus NS5 [30]. RdRp from HCV tolerated NaCl concentrations until 100 mM without significant RdRp activity loss [34]. However, it is unclear how Dengue and KFDV/Usutu NS5 proteins differ in their sensitivity to different concentrations of NaCl. Mechanistically, the binding of Dengue NS5 to ssRNA has been shown to release six Na^+^ ions [35]. Based on this report, the loss of activity can be speculated to be due to the dissociation of RNA at higher concentrations of NaCl and not due to the denaturation of protein. At a pH of 7.0, maximal activity was observed, tapered at pHs 5.0 and 9.0. The optimal activity was observed at a temperature of 30°C at pH 7, without significant activity loss at 42°C (Fig 2). Apparent K_m_,ATP was determined to be 42.15 ± 5.81 μM for KFDV NS5, under the tested conditions, as described in methods (Fig. 3). The reported values of k_m,GTP_ for glutathionylated Dengue and Zika NS5 were 189.8 ± 52.5 μM, and 137.2 ± 41.3 μM, respectively [36]. Motif F from the ring domain of RdRp with highly conserved Lys, Glu, Arg residues, and motif G from pinky finger domain, with conserved ala, binds to NTPs, and + 1/+2 junction of the template strand, respectively (Fig. 4) [37]. The comparable values of apparent K_m,ATP_ to K_m,GTP_ validates the use of the purified protein for inhibition studies.

**Fig. 2.**
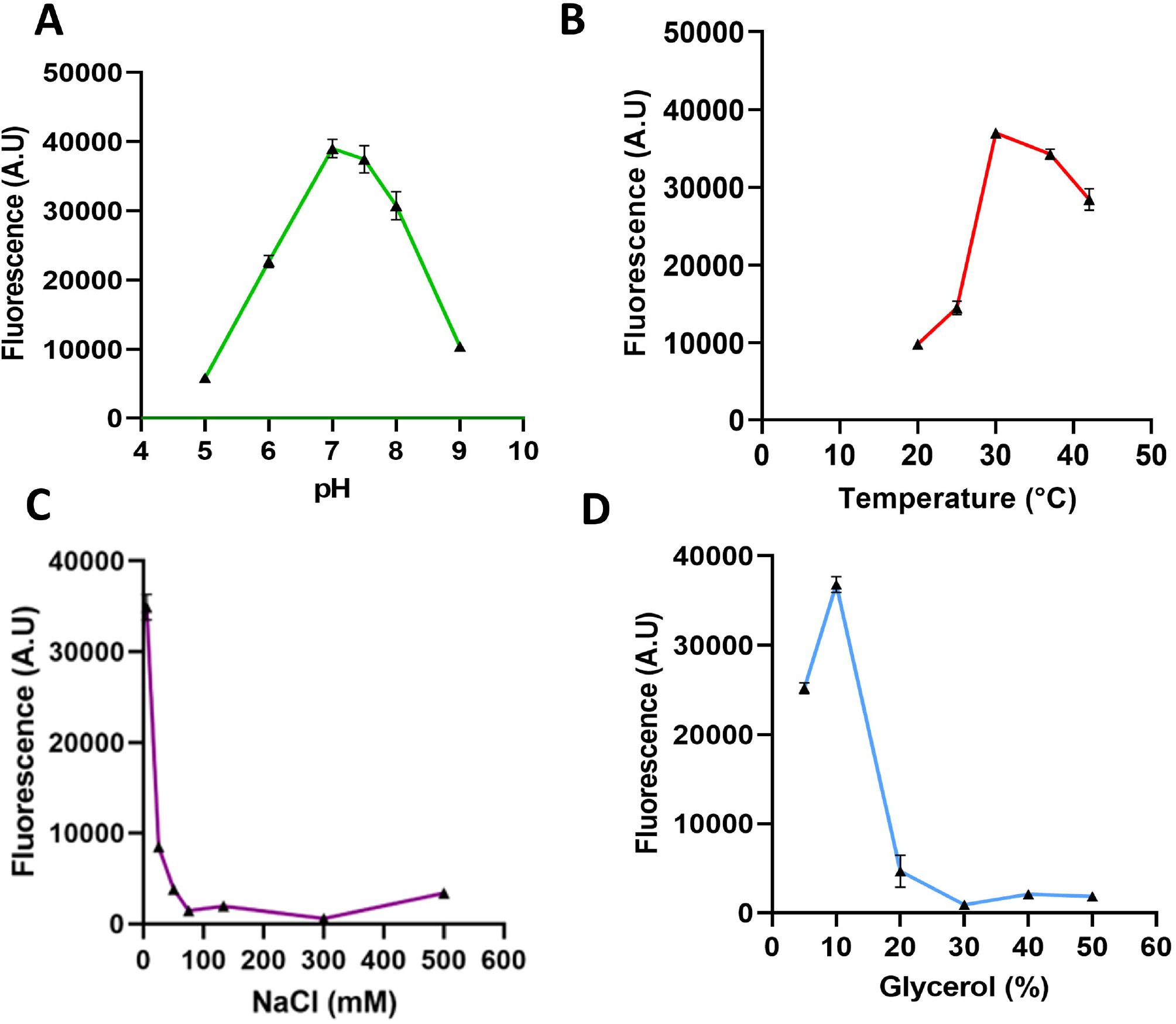
Optimization of reaction conditions for RdRp activity of KFDV NS5. Effect of pH (A), NaCl (B), glycerol (C), and temperature (D) on RdRp activity. Activity is represented as arbitrary fluorescence units (A.U.) on the Y-axis. For every sample well, fluorescence units were subtracted from a blank control devoid of NS5 protein under the same conditions. The standard deviation was computed and represented as error bars for every data point.

**Fig. 3.**
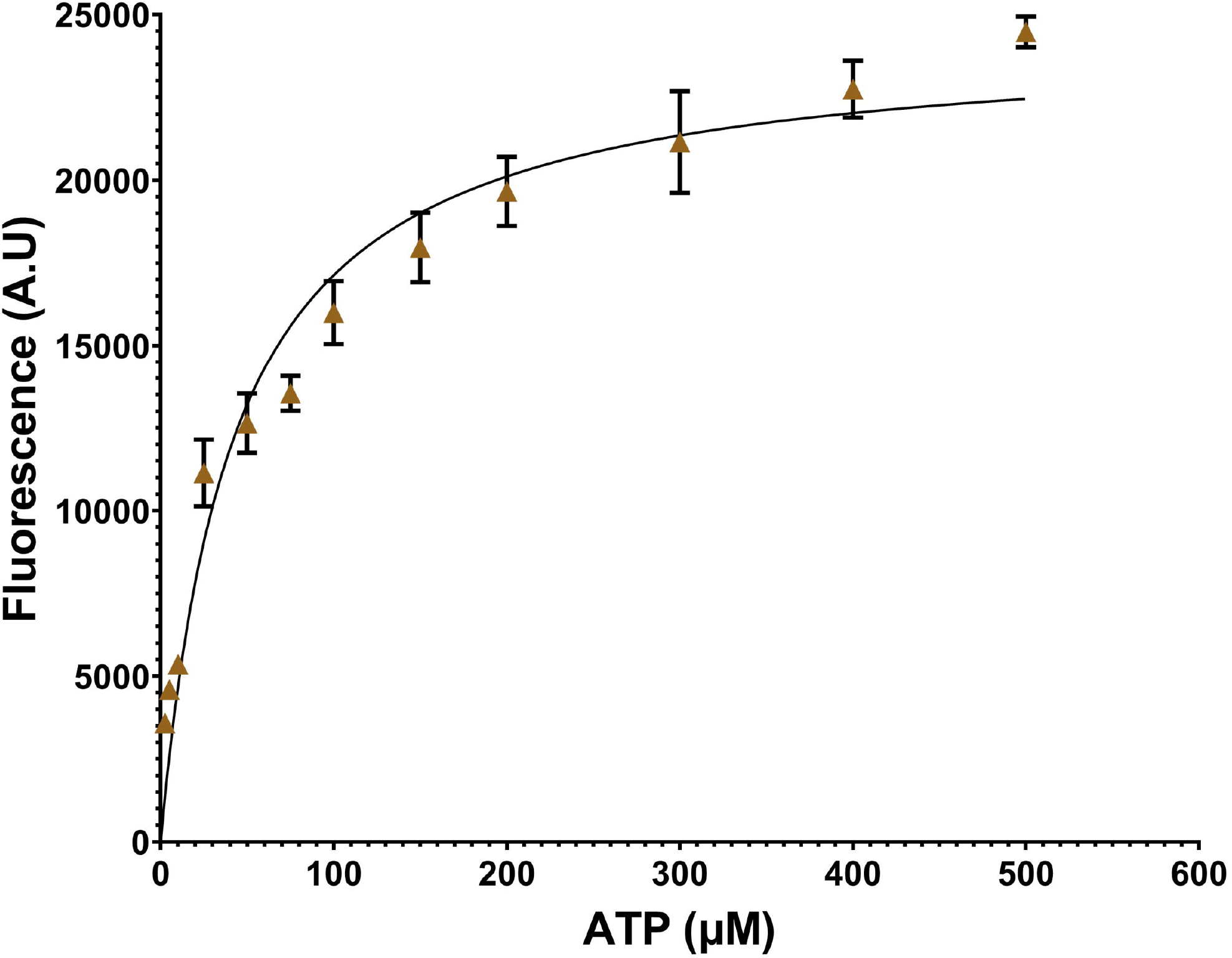
Determination of apparent K_m,ATP_ for RdRp domain of KFDV NS5. Apparent K_m_ for ATP was determined by Michaelis-Menten equation using graph pad prism 9.3.0. The representative graph contains values averaged from at least three independent experiments (n=3). Activity is represented as arbitrary fluorescence units (A.U.) on the Y-axis. The standard deviation error bars are shown for every data point.

**Fig. 4.**
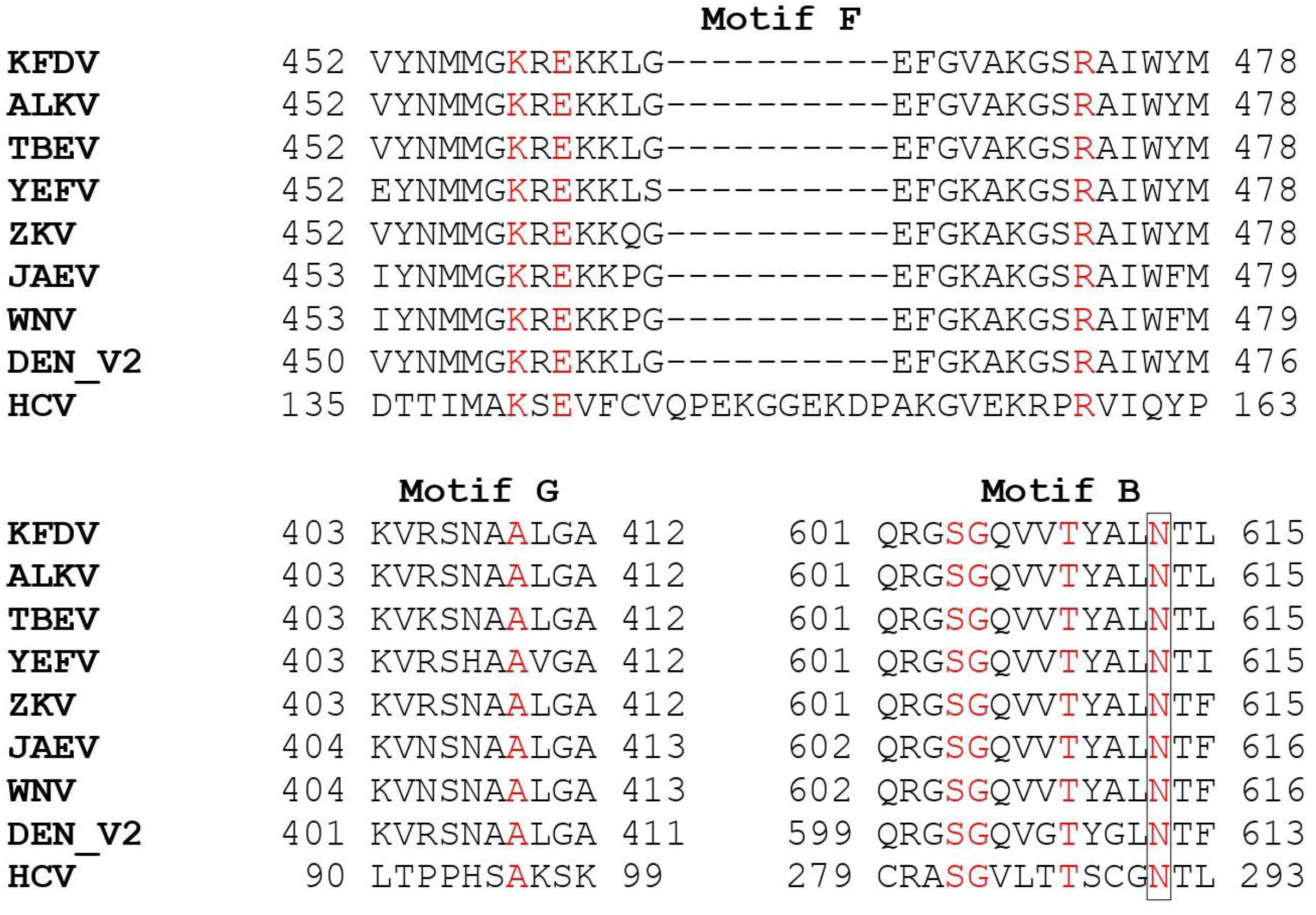
Amino acid sequence alignment of motifs F, G, and B of NS5 from the Flaviviridae family. Kyasanur forest disease virus (KFDV), Alkhurma hemorrhagic fever (ALKV), Tick-borne encephalitis virus (TBEV), Yellow fever virus (YEFV), Zika virus (ZKV), Japanese encephalitis virus (JAEV), West Nile virus (WNV) Dengue virus serotype 2 (DENV2), Hepatitis C virus (HCV). Conserved residues in Motif F and motif G, from ring and pinky finger domains involved in interacting with NTPs and the template, respectively, are highlighted. Asn291 from the NS5 RdRp domain of HCV makes a critical interaction with Sofosbuvir [33]. Asn291 is highly conserved across the sequences shown in the figure (marked as *). There is a higher level of conservation among the flavivirus genus than flaviviruses and HCV (which belong to the genus Hepacivirus).

### Sofosbuvir and sofosbuvir triphosphate inhibit the RDRP activity of KFDV NS5

With HCV NS5, the half-maximal inhibitory concentrations of sofosbuvir triphosphate and sofosbuvir were reported to be 0.12 μM and 2.06 ug/mL (corresponding to 3.8905 μM), respectively (24, 57). X-ray crystal structure of HCV NS5B with sofosbuvir (PDB: 4WTG) showed a critical interaction of Asn291 with the 2’-F group of Sofosbuvir, leading to its incorporation into the growing RNA chain and, subsequently, chain termination (Fig. 4) [39]. As indicated earlier, active site residues in the Flaviviridae family are highly conserved, including Asn291 increasing the likelihood of Sofosbuvir inhibiting KFDV NS5. Sofosbuvir and sofosbuvir triphosphate exhibited IC50 of 3.45 ± 0.012 and 3.73 ± 0.033 μM, respectively against KFDV NS5 (Fig. 5). Interestingly, Dasabuvir, a nonnucleoside analog, did not show inhibition of RdRp activity, even at 100 μM, under the tested conditions. IC_50_ of Dasabuvir varied between nanomolar and micromolar concentrations when tested against different genotypes of HCV RdRps [25]. Also, Dasabuvir is not predicted to bind to conserved residues in RdRps across other viruses [40].

**Fig. 5.**
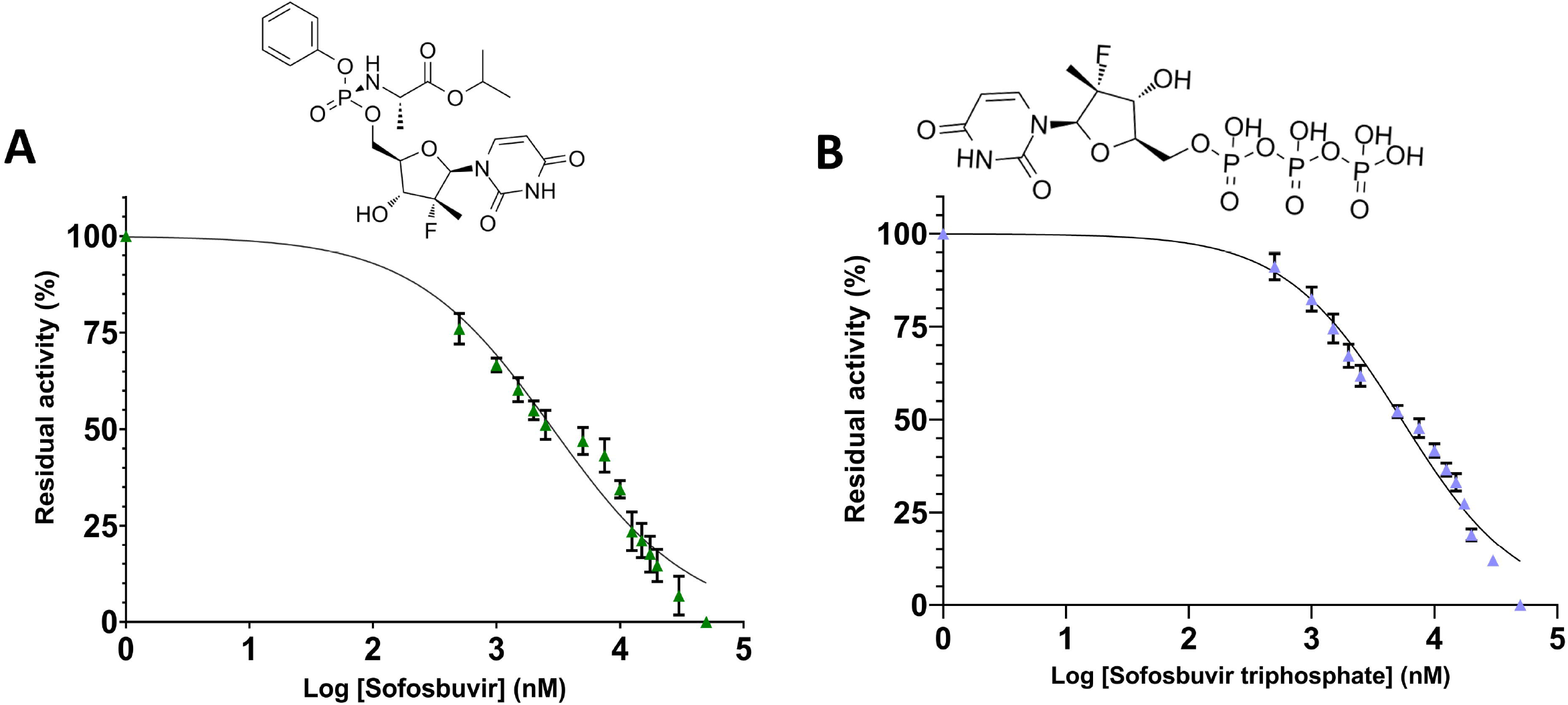
Determination of IC_50_ for sofosbuvir (A) and sofosbuvir triphosphate (B). Increasing concentrations of Sofosbuvir and sofosbuvir triphosphate were titrated using the optimized assay conditions described in the methods. The log of compound concentration is plotted against percentage activity and IC_50_, computed from the profiles using Graph pad prism 9.3.0 (n=3). The structures of compounds are shown above the IC50 graphs.

## Discussion

Kyasanur forest disease virus is a high-risk category pathogen endemic to southern India. The current line of treatment for KFD includes maintaining hydration, blood work, and management of neurological and hemorrhagic conditions [5, 6]. Sofosbuvir, also known as L-Alanine, N-[[P(S),2’R]-2’-deoxy-2’-fluoro-2’-methyl-P-phenyl-5’-uridylyl]-,1-methyl ethyl ester (C_22_H_29_FN_3_O_9_P), is an orally available nucleotide analog, approved for the treatment of HCV viral infections [26]. In preclinical studies, Sofosbuvir has exhibited inhibitory activity against all genotypes of HCV [41]. Sofosbuvir has been indicated for chronic HCV infection (genotype 1, 2, 3, or 4) in adults as part of combination therapy and in children older than 3 years with genotype for chronic HCV infections with ribavirin. Many Clinical trials are ongoing for evaluating sofosbuvir in different populations and dosing regimens in combination with other drugs against HCV [42 – 46]. Sofosbuvir also exhibited inhibitory activity against Dengue, Chikungunya, Yellow fever virus, Hepatitis E virus, and SARS-CoV2 [47 –51]. Sofosbuvir (PSI-7977) is metabolized to its triphosphate metabolite, β-D-2’-deoxy-2’-fluoro-2’-C-methyluridine 5’-triphosphate (PSI-7409), by hydrolysis of the carboxyl ester moiety, spontaneous release of the Phenyl group, deamidation and phosphorylation [52, 53]. KFDV NS5 expression construct used in this study also included the MTase domain, as the MTase was demonstrated to influence the activity RdRp domain by interaction with the template RNA [38]. KFDV NS5 protein was recombinantly expressed in bacteria, and so were NS5 proteins from Usutu, Zika, Japanese encephalitis, Dengue, HCV, and West Nile virus [30, 33, 37, 54, 55, 56]. Recombinantly purified NS5 RDRP protein from the Zika virus exhibited an IC_50_ of 0.38 ± 0.03 μM against sofosbuvir triphosphate [56, 47]. The inhibitory concentration (IC50) of sofosbuvir triphosphate and sofosbuvir were in the same range for KFDV NS5. This could be due to differences in the architecture of sofosbuvir binding sites across viruses. This is the first report on the expression and purification of the NS5 protein of KFDV, its assay establishment, and the demonstration of inhibition of RdRp activity by sofosbuvir. Further, the established biochemical assay can screen potential inhibitors of KFDV RdRp. Drug repurposing approaches against KFD, a neglected disease, can pave the way for identifying novel inhibitors for developing targeted therapeutic interventions against the disease.

## Abbreviations

KFD: Kyasanur forest disease
KFDV: Kyasanur forest disease virus
NS5: Non-structural protein 5
RdRp: RNA-dependent RNA polymerase
MTase: Methyl transferase
HCV: Hepatitis C virus

## Acknowledgments

The authors thank Tata Trusts for funding this study. We would like to acknowledge the NCBS-inStem-CCAMP mass spectrometry facility for the technical assistance provided for this study.

## Availability of data and materials

All the relevant data used to support the findings of this study are included in the article.

